# Parallelism in gene expression between foothill and alpine ecotypes in *Arabidopsis arenosa*

**DOI:** 10.1101/2020.03.05.977892

**Authors:** Guillaume Wos, Magdalena Bohutínská, Jana Nosková, Terezie Mandáková, Filip Kolář

**Affiliations:** Department of Botany, Charles University, 128 01 Prague, Czech Republic; Institute of Botany, The Czech Academy of Sciences, 252 43 Průhonice, Czech Republic; Central European Institute of Technology and Faculty of Science, Masaryk University, 625 00 Brno, Czech Republic; Institute of Botany, University of Innsbruck, AT-6020 Innsbruck, Austria

**Keywords:** Alpine adaptation, *Arabidopsis arenosa*, common garden experiment, gene-by-environment interaction, gene expression analysis, parallel evolution

## Abstract

Parallel adaptation results from independent evolution of similar traits between closely related lineages and allows testing to which extent evolution is repeatable. Parallel adaptation often involves similar gene expression changes but the identity of genes shaped by parallel selection and the causes of expression parallelism remains largely unknown. By comparing genomes and transcriptomes of four independent foothill–alpine population pairs across four treatments, we addressed genetic basis, plasticity and functional consequences of gene expression parallelism in alpine adaptation. Seeds of four population pairs of *Arabidopsis arenosa* from distinct mountain regions were raised under four treatments that differed in temperature and irradiance, factors varying strongly with elevation. Parallelism in gene expression was quantified by RNA-seq in leaves of young plants. By manipulating temperature and irradiance, we also tested for parallelism in plasticity (gene-by-environment interaction [GEI]). We found significant parallelism in differential gene expression across four independently recruited alpine ecotypes with an overrepresentation of genes involved in biotic stress response. In addition, we demonstrated significant parallelism in GEI indicating shared response to environmental variation in our foothill and alpine populations. Fraction of genes showing expression parallelism also encompassed genomic variants showing outlying differentiation, with greater enrichment of such variants in cis-regulatory elements. In summary, our results suggest frequent evolutionary repeatability in shaping expression difference associated with colonization of an alpine environment and support the hypothesis of an important role of genetic variation in cis-regulatory elements in gene expression parallelism.

## Introduction

Parallel evolution is independent evolution of similar adaptations between closely related organisms (Savolainen, Lascoux, & Merilä, 2013). When exposed to similar environments, populations or closely related lineages often evolve similar phenotypic, physiological or behavioural features. Such a convergence in functional traits was supported by many empirical examples, in animals (Blackledge & Gillespie, 2004; Velotta et al., 2017), plants (Reich, Walters, & Ellsworth, 1997) or bacteria (Fong, Joyce, & Palsson, 2005). Studies on convergence of functional traits found a common genetic basis and often involved similar genetic changes, especially in cases of low divergence between lineages (Conte, Arnegard, Peichel, & Schluter, 2012; Stern & Orgogozo, 2009), suggesting that some genes are more likely to be subject to selection during parallel evolution. In general, studying the genetic basis of parallel evolution in repeated environments provides strong evidence for the roles of natural selection and reveal genes or pathways particularly important in adaptation under certain changing conditions (Elmer & Meyer, 2011).

Evolution of parallel phenotypes often involve similar gene expression changes, indeed, parallelism at the level of gene expression is common to many organisms (Cooper, Rozen, & Lenski, 2003; Zhao, Wit, Svetec, & Begun, 2015), and may be a source of important adaptive convergence (Ferea, Botstein, Brown, & Rosenzweig, 1999). Therefore, gene expression analyses have great potential to detect genes involved in parallel adaptation and discover mechanisms governing evolutionary predictability at the genotype-environment interface (Stern & Orgogozo, 2009). Gene expression is driven by a complex interplay of genetic and environmental factors, yet their role in shaping expression parallelism is largely unknown. Modifications in gene sequences are often translated into constitutive differences in gene expression and these differences are generally mediated by mutations in cis-regulatory elements (Stern & Orgogozo, 2009; Wittkopp & Kalay, 2012). Mutations in cis-regulatory elements are in theory more likely to cause gene expression parallelism due to reduced negative pleiotropy as compared to mutations in coding sequences, however, empirical evidence of this is scarce (Wray, 2007). In addition, the transcriptome of a particular individual reflects also its actual surrounding environment. Such genotype-by-environment interactions may also change in a predictable way over parallel pairs, suggesting adaptive transcriptomic plasticity. Once again, however, the extent of such interaction in naturally replicated systems remains largely unknown (Zhao et al., 2015).

Parallelism in gene expression using whole transcriptome sequencing has been extensively studied in animals, in insects (Reed et al., 2011; Zhao et al., 2015) and fish species (Derome & Bernatchez, 2006; McGirr & Martin, 2018) in particular. In turn, such studies in plants remain scarce and generally focused only on expression patterns of particular genes or metabolic pathways (Chen et al., 2014; Des Marais & Rausher, 2010; Streisfeld & Rausher, 2009). Moreover, transcriptomic inquiries of parallel adaptation in both plants and animals were generally conducted using only two replicates parallelism (two pairs of populations/species from and outside of the selective environment). This, however limits our inference on the magnitude and evolutionary drivers of transcriptomic parallelism. Such questions could be efficiently addressed only by leveraging comparisons over multiple independent population pairs, as has been demonstrated using both genomic and phenotypic data (Bolnick et al., 2009; Stuart et al., 2017) yet not by gene expression analyses.

Here, we tested whether there is significant parallelism in gene expression by screening the entire transcriptomes of four independently formed alpine ecotypes and their foothill counterparts from distinct mountain regions and aimed to identify its underlying genetic basis and plasticity. Alpine environment constitutes an ideal system for investigating adaptive differentiation because it is associated with sharp environmental gradients and the island-like distribution of mountain regions promote parallel colonization from lower to higher elevations. Such a unique naturally replicated setup allows not only for identifying parallelism but also quantifying intraspecific variability in parallelism between each pair of regions. Firstly, we compared gene expression of four replicated alpine colonists of *Arabidopsis arenosa* (Brassicaceae) with corresponding foothill counterparts from the same region under the same treatment (differential expression) as well as across varying environmental conditions modelled by four treatments (gene-by-environment interaction [GEI], a measure of environmentally-induced plasticity) and revealed significant parallelism in both measures across nearly all pairs of regions. We further investigated its underlying genomic basis by leveraging genome-wide divergence scans of the same populations, and potential environmental consequences by enrichment analyses, asking specifically: 1) Does the overall direction and magnitude of gene expression difference between foothill and alpine ecotypes vary with changing environment and between the mountain regions? 2) Do the distinct alpine ecotypes independently adapted to high elevations use similar genetic pathways? 3) Do we also observe parallelism for genes showing significant gene-by-environment interaction suggesting shared response to environmental changes? 4) Are the genomic variants underlying the parallel expression candidates located rather in cis-regulatory elements than in coding elements?

## Methods

### Plant material

Seeds of 3 maternal plants of *A. arenosa* were collected in four mountain regions (termed region hereafter) across Central and Eastern Europe (Fig. S1): Niedere Tauern in the Austrian Alps (NT), Făgăraş Mountain in Southern Carpathians in Romania (FG) and Vysoké Tatry (VT) and Zapadné Tatry (ZT) Mountain in Western Carpathians in Slovakia. NT, FG and ZT are occupied by tetraploid populations and VT is occupied by diploid populations (Kolář et al., 2016). For each region, we collected seeds from one foothill (between 600 and 1000 meters a.s.l) and one alpine (between 1700 and 2200 meters a.s.l) ecotype. In *A. arenosa*, the two ecotypes are morphologically distinct with smaller size, cushion-like rosette, shorter stems and larger flowers in the alpine ecotype (Wos et al., 2019). Previous studies on *A. arenosa* showed that alpine stands in the NT, FG and ZT+VT regions have been colonized independently from the adjacent foothill populations, each corresponding to distinct genetic clusters (C. European, S. Carpathian and W. Carpathian, respectively; Monnahan et al., 2019). In case of ZT and VT, each region has been colonized by different ploidy level, however, the cytotypes are genetically closely related (Wos et al., 2019, see also the Discussion).

### Rearing conditions

To reduce potential maternal effects of the original localities, we first raised one generation of plants in growth chambers under constant conditions (21/18 °C, 16/8 hrs day/night, light ∼300 umol m^−2^ s^−1^) in pots filled with a mixture of peat and sand (ratio 2:3). Seeds collected from such first generation were then raised under four experimental treatments that varied in temperature and irradiance. Temperature and irradiance are two environmental parameters associated with elevation that clearly distinguished our foothill and alpine ecotypes (Fig. S1). Two levels per each factor were combined in a full-factorial design resulting in four treatments: “High temperature: High irradiance” (Ht:Hi); “High temperature: Low irradiance” (Ht:Li); “Low temperature: High irradiance” (Lt:Hi); “Low temperature: Low irradiance” (Lt:Li). The two intermediate treatments: “High temperature: Low irradiance” and “Low temperature: High irradiance”, were used to mimic conditions experienced by plants at low and high elevations respectively. The two extreme treatments “High temperature: High irradiance” and “Low temperature: Low irradiance” were used to test for an effect of a rise temperature or irradiance on gene expression.

Seeds were first stratified for one week (4°C, constant darkness) and then germinated. under 15 h light/9 h dark, light intensity 150 µmol m^−2^ s^−1^, 21 °C and relative humidity 50 %. After 20 days, seedlings were split into four separate growth chambers and exposed to one of the four treatments. For all treatments, day length was set to 16 h light/8 h dark. Temperature and irradiance were gradually changed to reach: 18°C/13°C day/night in high temperature treatments or 10°C/4°C day/night in low temperature treatments and from 280 to 980 µmol m^−2^ s^−1^ during the day in high irradiance treatments or from 50 to 200 µmol m^−2^ s^−1^ during the day in low irradiance treatments (see Table S1 for details). Temperatures were chosen to reflect average values during the growth period, from April to June (average temperature measured in the Austrian Alps at 600 meters = 18 °C, at 2000 meters = 10°C) and irradiance based on average values reported in a previous study of ecotypic differentiation of alpine plants spanning similar elevation range (Bertel, Buchner, Schönswetter, Frajman, & Neuner, 2016). Treatments were applied until plants reached the 14-leaf stage so that plant materials for RNA-seq were collected on plants of similar developmental stage.

### Sample collection and RNA extraction

Plant materials were collected between 51 and 86 days after plants were transferred in separate growth chambers depending on the treatment, following different growth rates in different treatments. For transcriptome analysis we randomly selected one individual per maternal line for each population (8 populations × 3 maternal lines × 1 individual × 4 treatments = 96 plants sequenced in total). For each plant, we collected the seventh rosette leaf at a similar time point for all treatments and leaf samples were immediately snap-frozen in liquid nitrogen. None of the plants had flowered at the time of collection. Total RNA was extracted using the NucleoSpin miRNA kit including a DNase treatment step (Macherey-Nagel, Düren, Germany) and assessed the purity and quantity of RNA with a Nanodrop 2000 spectrophotometer (Thermo Scientific, Wilmington, DE, USA) and RNA integrity with Agilent 2100 bioanalyzer (Agilent Technologies, Palo Alto, CA, USA).

### Library preparation, sequencing and data processing

Library was prepared using the Illumina TruSeq Stranded mRNA Kit (Illumina Catalog # RS-122-9004DOC). Specific TruSeq adapters were ligated on the cDNA for each individual. Individual sequencing was carried out on Illumina HiSeq 4000 (Illumina, San Diego, CA, USA) on four lanes (4 lanes × 24 individuals) using 150-bp paired-end reads. After sequencing, raw data were filtered to remove low-quality reads. The data are available in Sequence Read Archive (http://www.ebi.ac.uk/ena) under the accession PRJNA575330 (Wos et al., 2020a).

Quality of each individual library was checked using the software fastqc (http://www.bioinformatics.babraham.ac.uk/projects/fastqc/). Sequencing generated between 8 and 36 million reads per individual. Overrepresented sequences, which corresponded to the TruSeq adapter, were trimmed using cutadapt (Martin, 2011). Because of high similarities between *A. arenosa* and *Arabidopsis lyrata* genome, reads for each individual were aligned on the *A. lyrata* reference genome (Hu et al., 2011) as has been successfully applied in previous genomic and transcriptomic studies in *A. arenosa* (Baduel, Hunter, Yeola, & Bomblies, 2018; Monnahan et al., 2019; Yant et al., 2013) with the version-2 annotation (Rawat et al., 2015) using hisat2 v2.1.0 (Kim, Langmead, & Salzberg, 2017). The number of reads mapped on each gene was counted with featureCounts v1.6.3 (Liao, Smyth, & Shi, 2013), we kept only the uniquely mapped reads.

### Differential gene expression

Read counts were then analysed using the edgeR v3.12.0 package (Robinson, McCarthy, & Smyth, 2010) in R (R Core Team, 2013). We scaled the library size with the ‘calcNormFactors’ function, estimated dispersion using “estimateDisp” and obtained, for each region, list of differentially expressed genes between foothill and alpine ecotype using “glmFit” function. *P* values were adjusted for multiple testing with the Benjamini and Hochberg false discovery rate correction (FDR). Genes were considered as differentially expressed if FDR < 0.05 and as parallel if they were expressed in at least two regions and in the same direction.

We tested for the interaction ecotype × environment (GEI) as described in Levine, Eckert and Begun (2011), we performed three different contrasts using edgeR: 1) ecotype × alpine environment interaction as (foothill in Lt:Hi - foothill in Ht:Li) - (alpine in Lt:Hi - alpine in Ht:Li), 2) ecotype × temperature interaction as (foothill in Lt:Li - foothill in Ht:Li) - (alpine in Lt:Li - alpine in Ht:Li) to test for the effects of changes in temperature when irradiance was kept low, we run the same contrast under high irradiance, and 3) ecotype × irradiance interaction as (foothill in Lt:Li - foothill in Lt:Hi) - (alpine in Lt:Li - alpine in Lt:Hi) to test for the effects of changes in irradiance when temperature kept low, we run the same contrast under high temperature. In the context of GEI, genes were considered as parallel only if they overlapped across at least two mountain regions.

### Enrichment analysis

Gene expression analysis was complemented by a Gene Ontology (GO) term enrichment analysis for biological processes using GO term finder (Boyle et al., 2004). We used the *A. thaliana* annotation, using orthology definition provided in the *A. lyrata* version-2 annotation (Rawat et al., 2015). GO terms were considered significantly enriched if the FDR-adjusted *P* value was < 0.05. REVIGO tool was then used to group them based on semantic similarities (Supek, Bošnjak, Škunca, & Šmuc, 2011).

### SNP calling and candidate SNP detection

In order to investigate genomic underpinnings of our parallel expression candidates, we investigated their association with genomic SNPs exhibiting extreme differentiation between corresponding foothill-alpine population pair. We complemented our transcriptomes with whole genome re-sequencing data from different individuals of the same populations as was the origin of our experimental plants partly published previously (Monnahan et al., 2019), partly as new data available at Sequence Read Archives (project ID SUB6592572; Wos et al., 2020b). In total we retrieved genome-wide SNP variation of 8 individuals per each population, except for one population (alpine population from NT) with 7 individuals (Table S2). The raw sequences were processed and SNPs were called following Monnahan et al. (2019). Shortly, we used trimmomatic-0.36 (Bolger, Lohse, & Usadel, 2014) to remove adaptor sequences and low quality base pairs. Trimmed reads were mapped to reference genome *Arabidopsis lyrata* (Hu et al., 2011) in bwa-0.7.15 (Li & Durbin, 2009) with default settings. Duplicated reads were identified by picard-2.8.1 (Broad Institute 2018) and discarded together with reads that showed low mapping quality (< 25). We used GATK (v. 3.7) to call and filter reliable variants, following best practices (McKenna et al., 2010). Namely, we used HaplotypeCaller to call variants per individual with respect to its ploidy level and GenotypeGVCFs to aggregate variants for all samples. We selected only biallelic SNPs and removed those that matched following criteria: Quality by Depth (QD) < 2.0, FisherStrand (FS) > 60.0, RMSMappingQuality (MQ) < 40.0, MappingQualityRankSumTest (MQRS) < −12.5, ReadPosRankSum < −8.0, StrandOddsRatio (SOR) > 3.0. We further removed variants from sites with average read depth higher than 2 times standard deviation and regions with excessive heterozygosity indicating likely duplicated and paralogous regions respectively (for further details see Monnahan et al. (2019). In the final vcf, for each variant, we discarded genotypes with read depth lower than 8 ^×^ and with more than 20% genotypes missing.

We identified genome-wide differentiation single nucleotide polymorphism (SNP) outliers as 5% outliers from genome-wide distribution of F_ST_. We annotated each outlier SNP and assigned it to a gene, regulatory or intergenic variant using SnpEff 4.3 (Cingolani et al., 2012) following *A. lyrata* version-2 genome annotation (Rawat et al., 2015). We used standard terminology of SnpEff 4.3 software to define cis-regulatory elements and coding sequences, we considered all SNPs located in 5’ UTRs, introns, 3’ UTRs and in a 5 kb portion up- and down-stream of a gene as variation in cis-regulatory elements and all SNPs in exons as variants from coding sequences. We further restricted our candidate list to genes containing at least 3 SNP candidates to minimize the chance of identifying random allele frequency fluctuation rather than selective sweeps within a gene.

### Statistical analysis

We used permutational multivariate analysis of variance (PERMANOVA) to test for overall differentiation among the transcriptomic profiles (characterized by normalized read counts) categorized according to treatment, ecotype and region. We first created multidimensional scaling (MDS) plots using the ‘MDSplot’ function in edgeR v3.12.0 (Robinson et al., 2010) and extracted the corresponding distance matrix. The distance matrix was then used to compute PERMANOVA test (adonis2 function, vegan package; Oksanen et al., 2013), number of permutation = 10000) in RStudio (RStudio Team, 2015) using treatment, ecotype and their interaction or treatment, region and their interaction as predictors.

We quantified the degree of transcriptome-wide parallelism using a vector analysis as described in Stuart et al. (2017). We calculated difference in length and angle between each pair of regions (= proxy of parallelism) that are related to magnitude and direction of shift in gene expression. For each treatment, we calculated (1) the magnitude of divergence measured as the difference in vector lengths (Euclidian distance between foothill and corresponding alpine centroid from the same region) between pairs of regions and (2) the direction of divergence measured as the angle between any pairs of vectors. Then, we tested for the effects of region and treatment on magnitude and direction using linear model and visualized major trends in transcriptome similarity on MDS plots.

We tested for non-random overlap in gene expression candidate lists across mountain regions (i.e. significant parallelism) using Fischer’s exact test implemented in SuperExactTest package (Wang, Zhao, & Zhang, 2015).

In order to investigate genomic underpinnings of our parallel expression candidates, we tested for over-representation of genomic SNPs exhibiting extreme differentiation between corresponding foothill-alpine population pairs in different genomic regions associated with the parallel expression candidates (cis-regulatory elements vs. coding sequence). For each mountain region and genomic element category, we performed hypergeometric tests (*phyper* function; RStudio Team, 2015) to test for outlier SNPs enrichment in our lists of parallel expression candidates compared to the background of all divergence outliers (total number of 5 % outlier SNPs in cis-regulatory and coding elements for each mountain region).

In addition, for each pair of regions (six pairs in total) we identified parallel DEGs that harboured at least one SNP exhibiting excess alpine-foothill differentiation in both regions (i.e. parallel DEGs with underlying parallel outlier SNP). We then run hypergeometric tests (*phyper* function; RStudio Team, 2015) to test for significant enrichment of parallel outlier SNPs in cis-regulatory or in coding elements compared to the background (total number of 5 % outlier parallel SNPs in cis-regulatory and coding elements located at the same position for each pair of regions).

## Results

### General effects of treatment, ecotype and region on gene expression

All factors, i.e., treatment, mountain region and ecotype significantly affected gene expression in *A. arenosa*. MDS plot of the 96 samples (Fig. S2) and PERMANOVA test showed that treatment explained the highest proportion of variance in gene expression among the 96 samples (35.15 %, *F* = 16.62, *p* < 0.001), followed by region (20.24 %, *F* = 14.53, *p* < 0.001) and ecotype (5.20 %, *F* = 7.88, *p* < 0.001). Only the interaction term treatment × region was significant (7.48%, *F* = 1.79, p < 0.01) but not the interaction treatment × ecotype, suggesting that overall transcriptomic response of ecotypes was consistent across treatments. When taking individual environmental parameters separately, the effect of temperature was significant (10.25%, *F* = 10.74, *p* < 0.001), neither its interaction with region nor with ecotype was significant. Effect of irradiance was also significant and explained a greater proportion of the total variance than temperature (22.70%, *F* = 27.61, *p* < 0.001) and the interaction irradiance × region was also significant (3.86 %, *F* = 2.13, *p* < 0.05).

### Parallelism in gene expression between foothill and alpine ecotype

First, we quantified the degree of overall transcriptome-wide parallelism using a quantitative approach based on vector relationships among regions (corresponding pairs of alpine-foothill populations) in multidimensional space (Fig. 1, Table S3). Based on four replicates (treatments) we tested whether there was consistent response in the magnitude (difference in length of pairs of vectors) or direction (angle between pairs of vectors among the regions) of the transcriptomic parallelism between the regions. Transcriptome-wide parallelism in alpine-foothill gene expression was not consistent across treatments, as region had no significant effect neither on difference in length (ranging from |0.01| to |2.27|) nor angle (ranging from 10° to 149°).

**Fig. 1.**
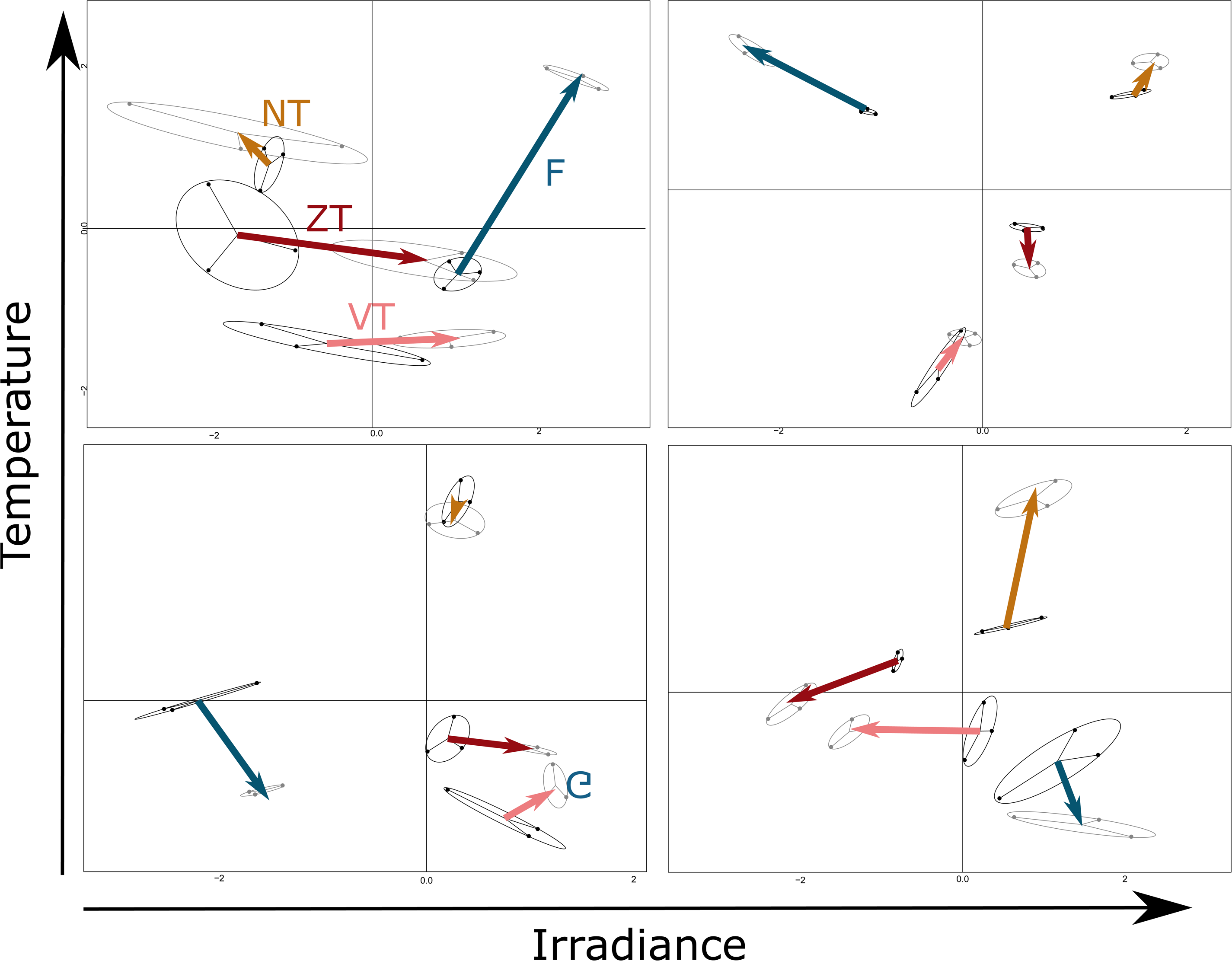
Multidimensional scaling (MDS) plots of gene expression data in each of the four treatments that vary in temperature (y-axis) and irradiance (x-axis) (*N* = 24 individuals per treatment). Individual plants were grouped by ecotype (black = foothill ecotype, grey = alpine). Arrows represent vectors connecting foothill and alpine centroid for each mountain region (NT = Niedere Tauern (Austria), FG = Făgăraş (Romania), VT = Vysoké Tatry (Slovakia), ZT = Západné Tatry (Slovakia)).

To quantify parallelism on a per-locus basis, we identified differentially-expressed genes (DEGs) between foothill and alpine populations for each region separately and then overlapped them, separately within each of the four treatments (i.e. a ‘common garden’ approach; Fig. 2, Table S4). Genes exhibiting parallel changes in expression (parallel DEGs) were those consistently up- or down-regulated in alpine populations in at least two regions. We revealed pervasive parallelism at the locus level, as the number of parallel DEGs in each intersection was significantly greater than expected by chance in all but one case; their total number ranged between 390 and 819 across treatments (Fig. 2, Table S5). We then run a GO term enrichment analysis to further identify the biological processes in which parallel DEGs were involved, based on all the genes showing parallel changes across the four treatments (*N* = 2179) to cover wide range of conditions in which the ecotypic effect could have been manifested. GO term enrichment revealed 182 GO terms significantly enriched at FDR < 0.05 grouped into 17 categories using REVIGO (Table S6). Most of the enriched GO terms felt into two categories: “response to other organisms” related to biotic stress response and “secondary metabolism”.

**Fig. 2.**
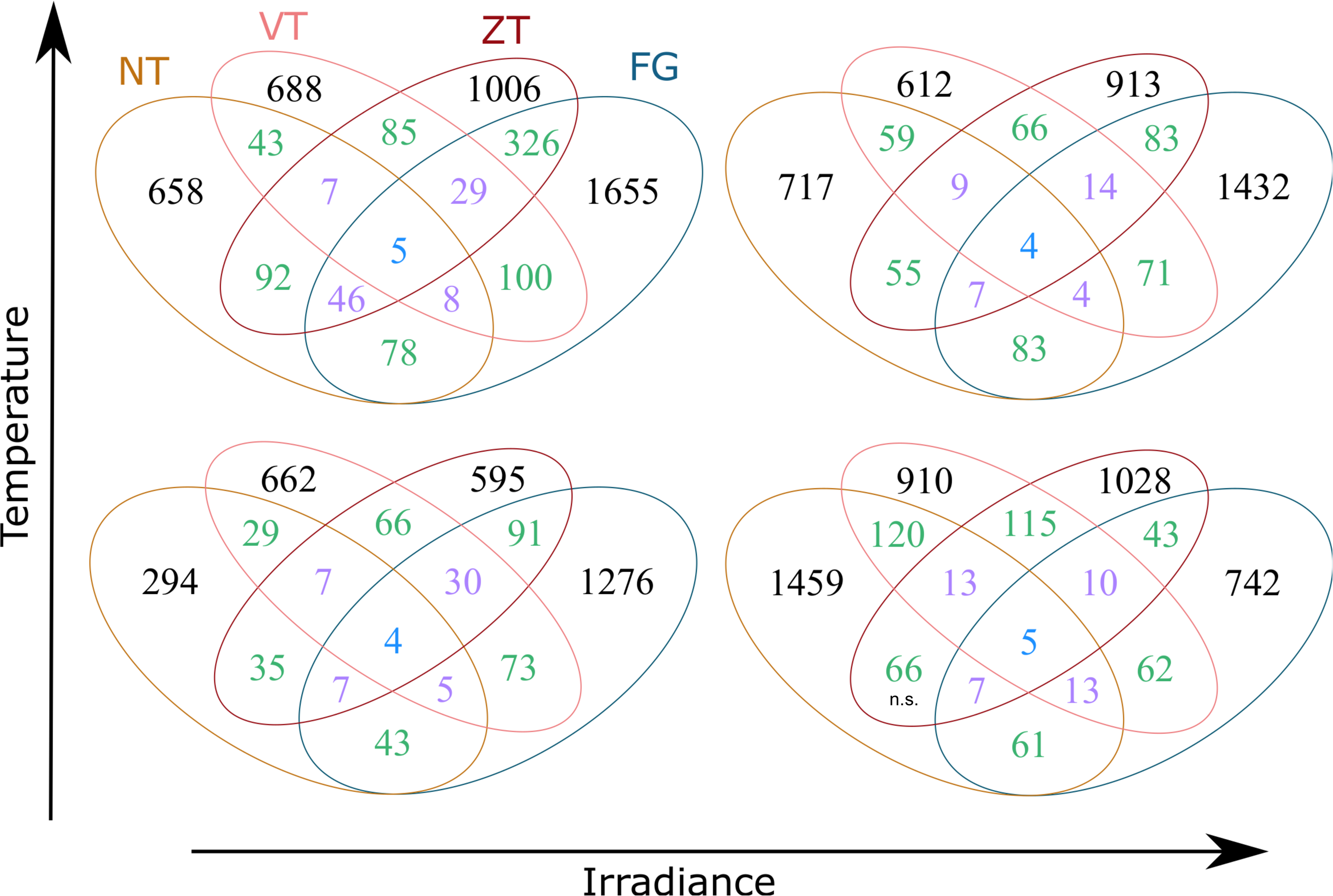
Number of differentially expressed genes (DEGs) between foothill and alpine ecotypes in each treatment. Colours depict the overlaps across two (green), three (purple) and four (blue) regions. NT = Niedere Tauern (Austria), FG = Făgăraş (Romania), VT = Vysoké Tatry (Slovakia), ZT = Západné Tatry (Slovakia). n.s. indicates non-significant overlap as identified by SuperExactTest.

We further focused on the strongest candidates for parallelism, i.e. genes showing overlap across all four regions in at least one treatment (10 genes in total, Table S5). Four of these genes had a known function in *A. thaliana*; three were related to defense response (*BETA-AMYLASE 5, FARNESOIC ACID CARBOXYL-O-METHYLTRANSFERASE*, LURP-one family) and were all down-regulated in alpine ecotypes, one gene was related to protein degradation (*F-BOX/LRR-REPEAT PROTEIN 25-RELATED*) and was up-regulated in alpine ecotypes. In the same way, DEGs that overlapped across three regions in at last one treatment (N = 181 in total, Table S5) were also mainly related to defense pathways and response to biotic stress (31 enriched GO terms of which 23 were related to biotic stress; data not shown).

### Genomic underpinnings of the parallel differential expression

Given the importance of mutations in cis-regulatory elements in parallel expression changes, we investigated whether genes showing parallelism in differential expression for each mountain region exhibited significant enrichment of highly differentiated SNPs (outliers for F_ST_ divergence between alpine vs. foothill population in that region) both in their coding sequence and in cis-regulatory elements (Table 2A). We found that parallel DEGs were significantly enriched for SNP differentiation outliers (as compared to the background of all divergence outliers) in cis-regulatory elements in two regions, VT and ZT. In turn, we did not find significant enrichment for outlier SNPs in coding elements in any region. We then performed pairwise comparison between the mountain regions (six pairs in total) to test whether parallel genes common to two regions harboured at least one SNP exhibiting excess alpine-foothill differentiation at the same position (i.e. parallel outlier SNP). For each pair of regions, we identified DEGs with shared underlying outlier SNPs but the number varied greatly, generally corresponding with biogeographic differentiation among the regions (lowest shared proportion was found in comparisons with the Alpine, NT, region while highest within the Carpathian Mts. (FG, VT, ZT, table S7). The number of parallel outlier SNPs was significantly enriched in cis-regulatory elements only in the NT-VT region pair and in coding elements in the FG-ZT region pair (hypergeometric test, Table S8). By merging all the parallel DEGs sharing at least one parallel outlier SNP for the six pairs of region, GO term enrichment and REVIGO clustering revealed that these genes were mainly related to “response to other organisms” (biotic stress, Table S7).

### Parallelism in plastic response of ecotypes

Finally, we tested for parallelism in differential response of originally alpine and foothill populations to changing environmental conditions simulated by varying temperature and irradiance. Firstly, we quantified the overall level of parallelism by magnitude and direction of the vectors characterizing individual regions (foothill-alpine pairs) (Fig. 1). Neither any of the two environmental parameters individually, nor their joint treatment effects significantly affected vector magnitude and direction indicating no obvious signs of parallelism at the whole transcriptomic level.

To quantify plasticity in transcriptomic response on a per-locus basis, we further calculated genotype-by-environment interaction (GEI) between ecotypes and different environmental conditions separately for each region and then examined parallelism in such measure of plasticity as the degree of the overlap of the genes exhibiting significant GEI (Table 1). We assessed GEI between ecotype and three types of biologically relevant environmental contrasts: (i) conditions characteristic for the foothill and alpine environment (Fig. 3), (ii) changes of temperature when irradiance was kept constant (under high and low irradiance) and (iii) changes of irradiance when temperature was kept constant (under high and low temperature). In all environmental contrasts we detected significant overlaps in genes exhibiting GEI across all pairs of mountain ranges as well as in majority of triple overlaps (Fig. S4) altogether demonstrating significant parallelism in genes showing plastic expression response to important environmental parameters differentiating foothill and alpine sites.

**Table 1.** Number of differentially expressed genes (FDR < 0.05) showing significant (A) ecotype × alpine environment (list of genes in Table S9), (B) ecotype × temperature under low and high irradiance (list of genes in Table S10) and (C) ecotype × irradiance under low and high temperature (list of genes in Table S11) interaction in each of the four mountain regions. The last column shows the number of parallel DEGs that overlapped across at least two mountain regions for each interaction.

| Interaction |  |  | Mountain Regions |  |  |  | Parallel DEGs |
| --- | --- | --- | --- | --- | --- | --- | --- |
|  |  |  | Niedere Tauern (NT) | Făgăraș (FG) | Vysoké Tatry (VT) | Západné Tatry (ZT) |  |
| (A) Ecotype × alpine env. |  | Nb DEGs | 191 | 348 | 163 | 103 | 54 |
|  |  | F-value (R<sup>2</sup>) | **4.05\*\*\***<br>(21.6%) | **4.29\*\*\***<br>(20.2%) | **3.21\*\*\***<br>(19.2%) | **4.57\*\*\***<br>(20.2%) |  |
| (B) Ecotype × temperature | Under low irradiance | Nb DEGs | 132 | 372 | 221 | 394 | 118 |
|  |  | F-value (R<sup>2</sup>) | **3.37\*\*\***<br>(20.2%) | **6.54\*\*\***<br>(24.4%) | **3.85\*\*\***<br>(20.9%) | **5.70\*\*\***<br>(25.2%) |  |
|  | Under high irradiance | Nb DEGs | 109 | 50 | 103 | 138 | 21 |
|  |  | F-value (R<sup>2</sup>) | **4.63\*\*\***<br>(22.7%) | **3.73\*\***<br>(17.8%) | **2.93\*\*\***<br>(18.7%) | **3.89\*\*\***<br>(20.3%) |  |
| (C) Ecotype × irradiance | Under high temperature | Nb DEGs | 122 | 958 | 153 | 240 | 147 |
|  |  | F-value (R<sup>2</sup>) | **4.13\*\*\***<br>(21.4%) | **5.54\*\*\***<br>(20.4%) | **3.18\*\*\***<br>(17.7%) | **4.44\*\*\***<br>(15.2%) |  |
|  | Under low temperature | Nb DEGs | 207 | 85 | 140 | 356 | 66 |
|  |  | F-value (R<sup>2</sup>) | **4.34\*\*\***<br>(17.5%) | **3.10\*\*\***<br>(16.9%) | **3.63\*\*\***<br>(17.8%) | **4.46\*\*\***<br>(24.3%) |  |

F-value for the ecotype × treatment interaction for each region tested by PERMANOVA test and percentage of variance explained (R<sup>2</sup>) in parentheses are shown. \*\*\* $P < 0.001$ , \*\* $P < 0.01$

**Fig. 3.**
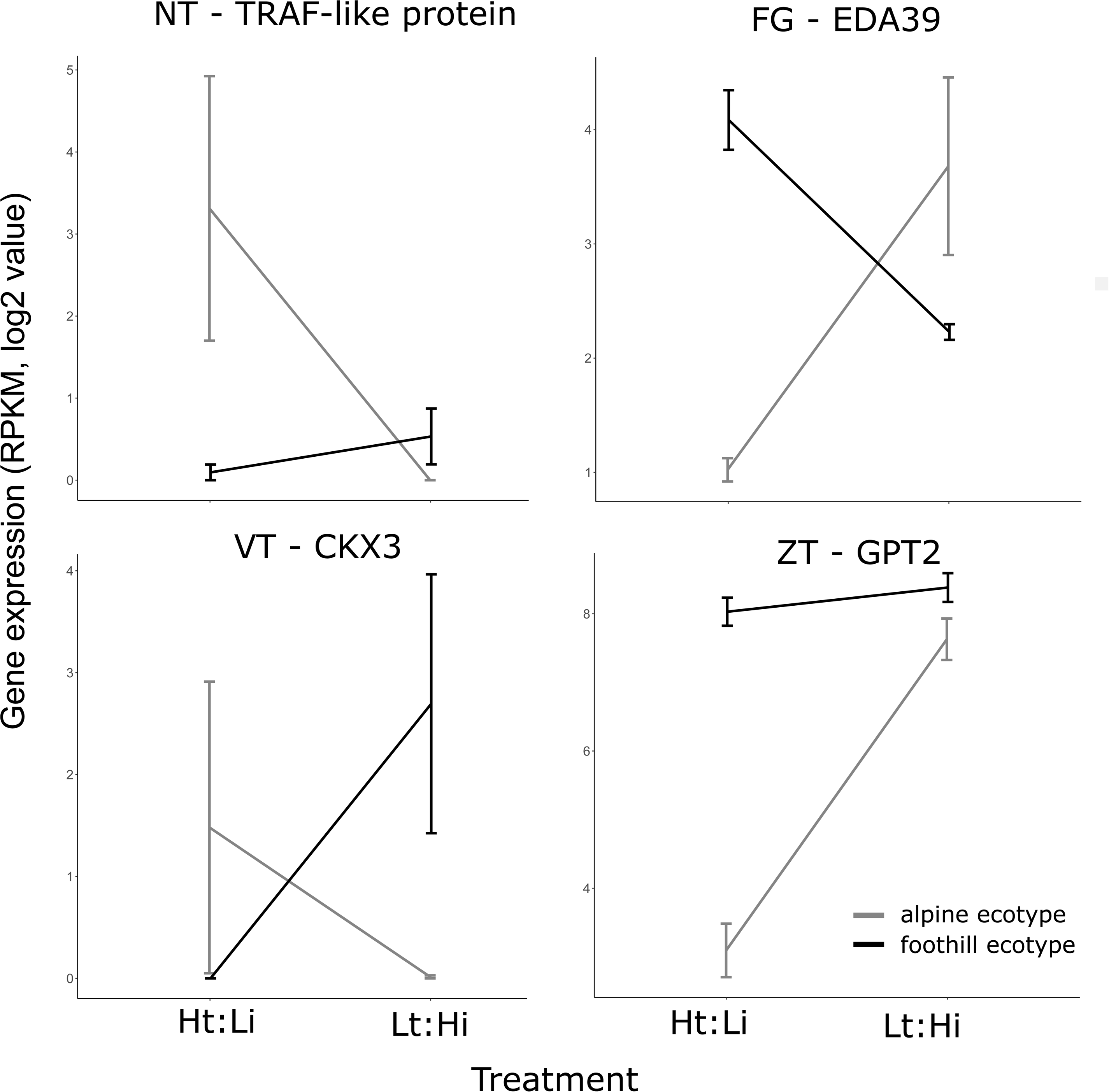
Examples of reaction norm of the most significant genes (lowest FDR-adjusted *P* value) with a known function showing ecotype × alpine environment interaction for each mountain region. Plot shows gene expression values (Reads Per Kilobase per Million mapped read - RPKM) for foothill (in black) and alpine ecotype (in grey) raised in “High Temperature: Low irradiance (Ht:Li)” (=approximating the low-elevation conditions) and “Low Temperature: High irradiance (Lt:Hi)” (=approximating the high-elevation conditions). NT = Niedere Tauern (Austria), FG = Făgăraş (Romania), VT = Vysoké Tatry (Slovakia), ZT = Západné Tatry (Slovakia). Abbreviations and gene function are: TRAF-like protein (Tumor necrosis factor receptor-associated factor - Immune system), EDA39 (*EMBRYO SAC DEVELOPMENT ARREST 39* - stomatal movement), CKX3 (*CYTOKININ OXIDASE 3* - cytokinin degradation) and GPT2 (*GLUCOSE-6-PHOSPHATE/PHOSPHATE TRANSLOCATOR 2* - regulation of photosynthesis).

Then, we examined potential function of such parallel GEI candidates. For treatment contrast approximating foothill vs. alpine conditions, enrichment analysis revealed five enriched GO terms (based on 54 such candidates; Table S9), four of them were related to cell wall modification (“hemicellulose metabolic process”, “cell wall polysaccharide metabolic process”, “xyloglucan metabolic process”, “cell wall macromolecule metabolic process”). The last GO term enriched was “response to chemical” and included 11 genes in response to various hormones.

For temperature, we identified 21 (under high irradiance) and 118 (under low irradiance) parallel GEI (Table S10). GO term enrichment analysis on all the genes affected by changes in temperature (21+118 = 139) revealed 56 enriched GO terms grouped into 7 categories by REVIGO mainly related to the response of temperature stimulus, flavonoid biosynthesis and secondary metabolism.

Finally, for irradiance, we identified 66 (under low temperature) and 147 (under high temperature) genes with significant ecotype × irradiance in parallel across regions (Table S11). GO term enrichment on all the genes (66 + 147 = 213) revealed 73 associated enriched GO terms. REVIGO grouped them into 10 categories mainly related to the response to other organisms and to the immune system.

Finally, we investigated whether the parallel genes showing GEI (all contrasts merged together) exhibit enrichment of differentiation outlier SNPs in their underlying genomic elements (Table 2B). We found significant enrichment for outlier SNPs in cis-regulatory elements in NT and VT regions, and only marginally significant in ZT, but an enrichment of SNPs in coding elements in FG region.

**Table 2.** Association between parallel expression candidates and genomic variation (outlier SNPs for alpine-foothill differentiation) in coding and associated cis-regulatory elements. (A) Genes showing parallelism in differential expression between the ecotypes and (B) genes showing parallel GEI.

| **A.** |  |  |  |  |  |  |  |
| --- | --- | --- | --- | --- | --- | --- | --- |
| **Regions** | **Nb genes** | **Cis-regulatory elements** |  |  | **Coding elements** |  |  |
|  |  | **Nb candidate SNPs** | **Total N of outlier SNPs** | **Enrichment** | **Nb candidate SNPs** | **Total N of outlier SNPs** | **Enrichment** |
| NT | 53 | 149 | 14490 | 0.79 | 98 | 8675 | 0.25 |
| FG | 263 | 1014 | 43078 | 0.71 | 831 | 34430 | 0.30 |
| VT | 290 | 1817 | 82330 | **0.03\*** | 1159 | 56511 | 0.98 |
| ZT | 618 | 4172 | 129164 | **0.02\*** | 2436 | 79539 | 0.98 |
| **B.** |  |  |  |  |  |  |  |
| **Regions** | **Nb genes** | **Cis-regulatory elements** |  |  | **Coding elements** |  |  |
|  |  | **Nb candidate SNPs** | **Total N of outlier SNPs** | **Enrichment** | **Nb candidate SNPs** | **Total N of outlier SNPs** | **Enrichment** |
| NT | 11 | 44 | 14490 | **< 0.001\*\*\*** | 8 | 8675 | 0.99 |
| FG | 75 | 243 | 43078 | 0.97 | 229 | 34430 | **0.04\*** |
| VT | 74 | 533 | 82330 | **< 0.001\*\*\*** | 214 | 56511 | 1.00 |
| ZT | 154 | 1053 | 129164 | 0.06(*) | 597 | 79539 | 0.95 |

The table shows (A) number of genes showing parallelism (within at least one treatment) and (B) number of genes showing parallel GEI (within at least one of the focal contrasts) for each mountain region with at least three candidate SNPs (at 5 % outliers), number of SNPs in cis-regulatory elements and coding elements and enrichment p-value (enrichment against the total number of SNPs (at 5 % outliers) for each region tested by hypergeometric test). \*\*\* $P < 0.001$ , \* $P < 0.05$

## Discussion

### Parallelism in gene expression between foothill and alpine ecotype and its genetic underpinnings

Taking advantage from a unique natural setup of four independently formed alpine ecotypes, we investigated intraspecific variation in parallelism in differential gene expression in a wild *Arabidopsis* species. Despite lack of consistent transcriptome-wide signal of parallelism in terms of magnitude and direction, we found significant overlap in differentially expressed genes (DEGs) between foothill and alpine ecotype across the four mountain regions, indicating pervasive significant parallelism at the level of individual sets of genes. However, the strength of association between parallelism in expression and its genomic underpinnings varied across regions and genomic elements. Specifically, significant enrichment of candidate SNPs (exhibiting excessive alpine-foothill differentiation, i.e., candidates for directional selection), was detected only in two of the four regions and exclusively in cis-regulatory elements. Pairwise comparisons of mountain regions revealed that candidate parallel SNPs (identical SNPs found as candidates in several regions) were significantly enriched in cis-regulatory elements in one pair of regions (NT-VT), and in coding elements in another regional pair (FG-ZT). In light of these results, cis-variants appear as potentially important players in shaping parallelism in gene expression, as opposed to variation in coding sequences that had mainly non-significant effect in our analyses. However, the lack for shared underlying SNPs across nearly all pairs of regions suggests that similar expression changes may be achieved rather by different mutations in cis-regulatory elements (Wittkopp & Kalay, 2012) and/or mutations in trans-acting regulatory elements (e.g., transcription factors) that were not addressable by our study design. Another explanation may involve varying patterns of directional selection between the different mountain regions.

Parallelism at the gene expression level, especially when replicated over more than two environmental transitions serves as a strong indicator of the role of natural selection in adaptive response to environmental triggers (Elmer & Meyer, 2011). Although convergent evolution is not always indicative of adaptive evolution (Losos, 2011), our case put forward the selective scenario as (i) all alpine populations occupy similar selective environment, (ii) significant overlaps in DEGs were found not only between pairs of regions but also across three or four geographically isolated regions rendering chance very unlikely and (iii) probably a minor contribution of genetic correlations or epistatic interactions in driving such relationships based on the high number of different metabolic pathways (182 enriched GO terms) in which parallel DEGs have been involved. Indeed genetic correlations are more likely to occur between genes of the same pathway (Phillips, 2008). While detailed follow-up studies are necessary for validation of the adaptive effect of the individual genes, our study brings one of the first transcriptome-wide evidence for pervasive parallelism from plants.

Interestingly, genes related to biotic stress response, to fungus and bacteria in particular, showed a strong and consistent signal of parallelism both in terms of gene expression (differential expression in the same direction) and SNP variation in the underlying genomic regions (differentiation outlier SNPs at the same position between pairs of regions). Elevation is associated with sharp variations in abiotic and biotic stress factors (Brown, Stevens, & Kaufman, 1996; Vetaas, 2002). In line, some studies identified pathways or metabolites related to biotic and abiotic stress associated with an elevational gradient. For instance, antioxidants (glutathione and phenol compounds) or defense proteins (Ma et al., 2015; Wildi & Lütz, 1996) varied strongly with elevation. In general, importance of abiotic factors increased (Körner, 2003) while importance of biotic factors decreased with elevation (Desprez-Loustau et al., 2010; Rasmann, Pellissier, Defossez, Jactel, & Kunstler, 2014). These results were consistent with our findings in particular that the overall decrease in pathogen and/or competition pressures with elevation was constitutively manifested at the gene expression level.

Despite original sites of the foothill and alpine ecotypes strongly differ in climatic conditions, we found only few GO terms directly and consistently associated with abiotic stress across regions among parallel candidates. Instead, we found some GO terms associated with general processes such as “metabolism” or “biological regulation” containing genes involved in responses to multiple environmental stresses and controlling many developmental aspects from germination to flowering. On one hand, these enriched GO terms may just reflect major changes that occur along an elevational gradient such as reduced partial pressure of CO_2_ with elevation that affects photosynthesis and the underlying metabolism (Wang et al., 2017). On the other hand, they may be also linked with the observed phenotypic convergence in *A. arenosa* alpine ecotypes in these mountain regions well-reflecting the abiotic pressures, i.e. reduced size or shifted flowering time (Měsíček & Goliašová, 2002). Additionally, we also cannot exclude that such major phenotypic differences may be due to selection on only few genes of larger effects that could be missed by enrichment analyses (e.g. *FRIGIDA* gene controlling flowering along a latitudinal gradient in *A. thaliana*, (Stinchcombe et al., 2004). However, genes controlling developmental aspects were lacking among our strongest parallelism candidates (DEGs overlapping three or four regions) ruling out a hypothesis of a single major-effect gene that would stand consistently behind the response to abiotic triggers.

Apart from parallelism in directional selection, similarity in gene expression may reflect historical events such as gene flow and effect of genetic drift. While the effect of genetic drift is likely minimized by absence of severe bottlenecks in *A. arenosa* populations (high and constant values of synonymous diversity and Tajima’s D across the species range, disregarding the ecotypes; Monnahan et al., 2019), previous study has detected complex reticulated evolution of diploid and tetraploid alpine ecotypes in from the Tatry Mountain, VT and ZT. However, gene flow had not markedly affected our measures of the magnitude of parallelism as the expression similarity among VT and ZT regions in terms of number of parallel DEGs fell well within the range of DEG overlap among all the other mountain regions. Similarly, there was no apparent effect of ploidy on parallelism as we observed similar overlap between regions occupied by tetraploids as was their overlap with the VT region represented by diploids. However, VT and ZT harbored more candidate SNPs at the same position than any other pairs of regions what may indicate overall sharing of genetic variation due to weak lineage sorting and/or recent gene flow.

### Parallelism in plastic response of ecotypes

Parallel adaptation does not have to be manifested only at level of differential expression *per se*. Parallel adaptive candidates may also exhibit a plastic response to important environmental triggers, that is manifested as gene-by-environment interaction (GEI) between adapted ecotype and manipulated environmental parameters. We detected significant GEI in response to two prominent abiotic factors affecting plant life in high elevations indicating that foothill and alpine ecotypes responded differently to environmental changes. Importantly, we detected significant parallelism in genes exhibiting such GEI across most of the regions although such overlap was considerably lower in absolute terms than parallelism at the level of DEGs (lower number of overlapping candidates and absence of complete, i.e. four-fold, overlaps). Parallel DEGs showing GEI tended to show greater enrichment for candidate SNPs in cis-regulatory elements in all but one region suggesting an important contribution of genetic variation in cis-regulatory elements in triggering similar expression changes in response to environmental stress.

In the context of ecotype × alpine environment, parallel DEGs showing significant GEI were hormone-related genes and cell-wall-modification enzymes, belonging to the XTH family involved in cell wall strengthening (Cosgrove, 2005). In plants, cell wall is one of the first mechanical barriers against abiotic and biotic stress and cell wall thickness was positively correlated with elevation in different species (Kogami, Hanba, Kibe, Terashima, & Masuzawa, 2001; Ma et al., 2015). When environmental variables, temperature and irradiance, were analyzed separately DEGs showing significant GEI were mainly related to response to temperature stimulus and plant defense respectively. The response to temperature stimulus includes many GO terms related to abiotic (temperature, oxidative and drought) factors known to vary along an elevational gradient (Ma et al., 2015). The overrepresentation of plant defense genes in response to changes in irradiance may be due to interactions between the light sensing and plant defense pathway (Karpinski, Gabrys, Mateo, Karpinska, & Mullineaux, 2003).

Local adaptation is commonly invoked to explain the maintenance of plasticity between populations (Josephs, 2018). In our system, based on the relatively high number of genes showing GEI, it is likely that foothill and alpine habitats differ in their fitness optima. Our results provided sets of potential candidate genes important for local adaptation. However, while changes in abiotic- and biotic-stress signaling compounds may be linked to an increased fitness in alpine environments, this requires experimental validation as does the overall quantification to what extent all the remaining non-parallel genes we identified contribute to plant fitness in an alpine environment.

## Conclusion

Our design involving comparison of transcriptomic and genomic profiles across four pairs of foothill and alpine ecotypes revealed on one hand pervasive parallelism at the “constitutive” level highlighting the prominent role of natural selection acting on genes and pathways related to biotic stress. On the other hand, we also demonstrated lower, yet still significant levels of parallelism at the “plastic” level indicating that populations of different origins may also exhibit shared responses to variation in the same environmental factors. In sum, our study demonstrates that the repeatability of evolution can be manifested at various levels of the complex genotype-environmental interface and that analysis of multiple replicated instances of ecotypic differentiation may aid uncovering such processes in natural populations.

## Acknowledgments

We thank Adam Knotek for his help with collecting plant material for RNAseq. Plant Sciences Core Facility of CEITEC MU is acknowledged for the cultivation of experimental plants used in this paper. Library construction and sequencing were performed at the Norwegian Sequencing Centre, University of Oslo. This work was supported by the Czech Science Foundation (project 17-20357Y to F.K.). Additional support was provided by the Norwegian Research Council (FRIPRO mobility project 262033 to F.K.), the Czech Science Foundation (17-13029S to T.M.), the CEITEC 2020 project (grant no. LQ1601 to T.M.) and by Ministry of Education Youth and Sports of the Czech Republic (7AMB18AT022 to G.W.). Computational resources were provided by the CESNET LM2015042 and the CERIT Scientific Cloud LM2015085, under the program Projects of Large Research, Development, and Innovations Infrastructures. The authors declare no conflicts of interest.

## Author contributions

GW and FK conceived and designed the study, GW and JN collected the data, GW and MB analyzed the data, GW and FK drafted the initial version of the manuscript and all authors contributed to later versions of the manuscript.

## Data accessibility

RNA-seq data used in this study are available at Sequence Read Archives (project ID PRJNA575330). We used a subset of the genomic data available at Sequence Read Archives (project ID SUB6592572) and from a previous study (project ID PRJNA484107; Monnahan et al., 2019) (details on the samples used for genomic data are available in Table S2).

