## Supplementary figures and images for "Parallelism in gene expression between foothill and alpine ecotypes in *Arabidopsis arenosa*"

### Fig. S4

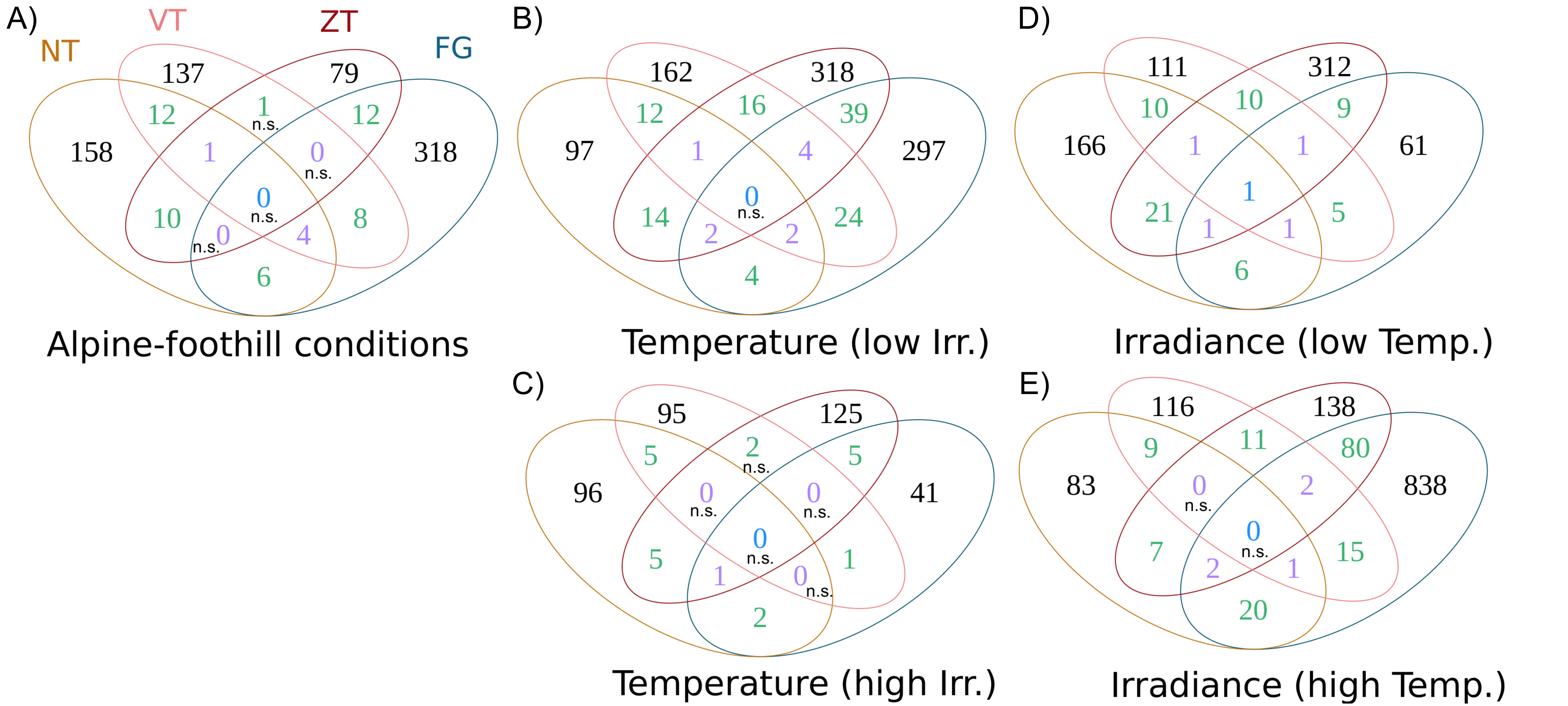
