## Supplementary material for "Parallelism in gene expression between foothill and alpine ecotypes in *Arabidopsis arenosa*": Fig. S1; Fig. S2; Table S8

### Supplementary information

Separate files:

**Fig. S4**. Numbers of differentially expressed genes showing significant (A) ecotype × alpine environment, (B and C) ecotype × temperature under low and high irradiance and (D and E) ecotype × irradiance under low and high temperature interaction. Colours depict the overlaps across two (green), three (purple) and four (blue) regions. NT = Niedere Tauern (Austria), FG = Făgăraș (Romania), VT = Vysoké Tatry (Slovakia), ZT = Západné Tatry (Slovakia). n.s. indicates non-significant overlap as identified by SuperExactTest.

**Table S1**. Rearing conditions of *Arabidopsis arenosa* used in this study

**Table S2**. Details on samples used for the genomic analysis

**Table S3**. Raw data and average values for magnitude and direction of vectors presented in Fig. 1.

**Table S4**. List of differentially expressed genes between foothill and alpine ecotypes for each mountain region and for each treatment

**Table S5**. List of differentially expressed genes that overlapped across at least two mountain regions for each treatment

**Table S6**. GO term enrichment and REVIGO clustering of parallel genes.

**Table S7**. Number of SNP at the same position for each pair of regions

**Table S9**. List of differentially expressed genes showing significant ecotype x alpine environment interaction, list of genes that overlapped across at least two mountain regions and GO term enrichment analysis on overlapping genes.

**Table S10**. List of differentially expressed genes showing significant ecotype x temperature interaction, list of genes that overlapped across at least two mountain regions, GO term enrichment analysis on overlapping genes and REVIGO clustering.

**Table S11**. List of differentially expressed genes showing significant ecotype x irradiance interaction, list of genes that overlapped across at least two mountain regions, GO term enrichment analysis on overlapping genes and REVIGO clustering.

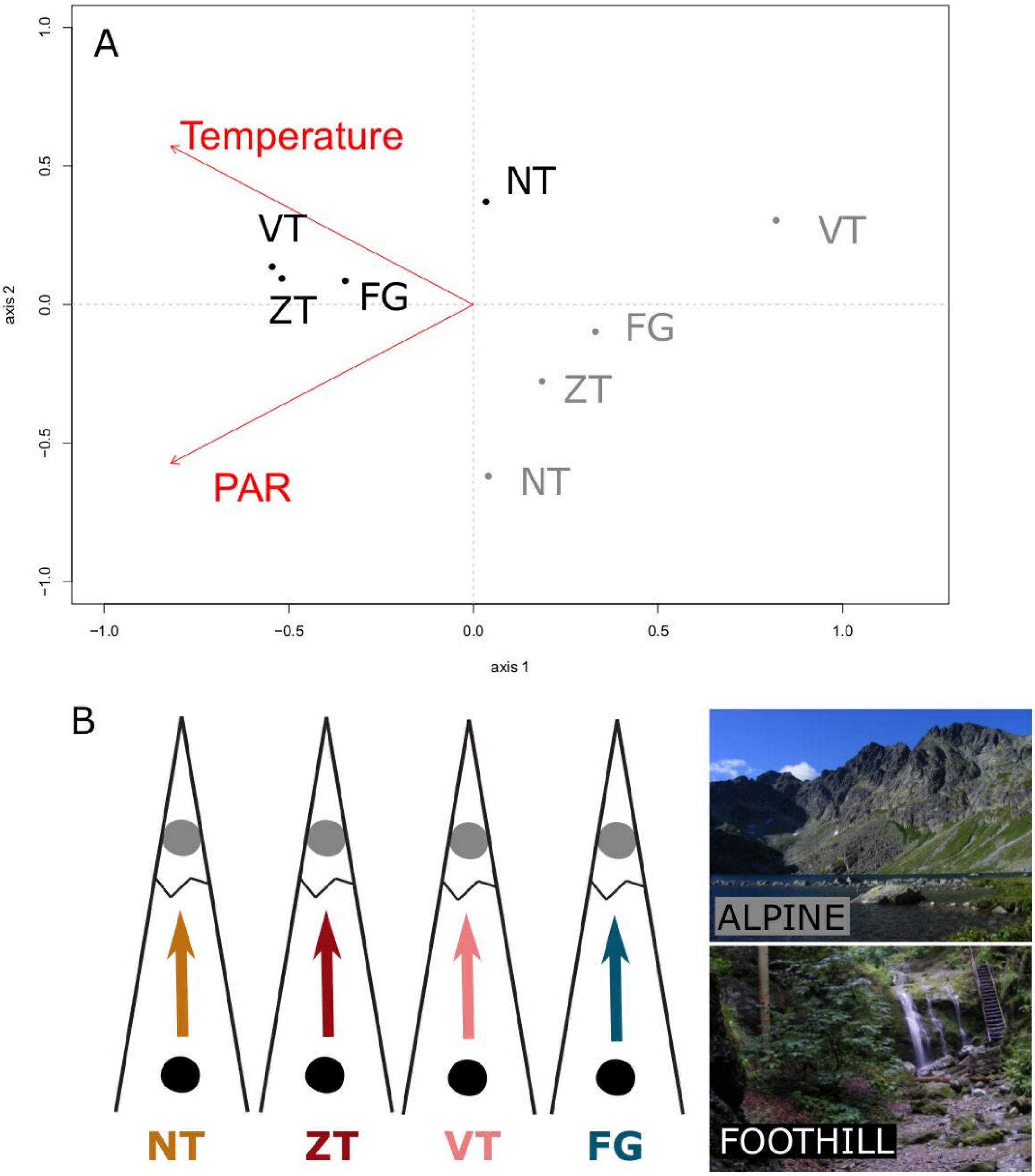


**Fig. S1**. A) Principal component analysis of the two parameters, temperature and irradiance, describing the environmental conditions at each native site of the 8 *A. arenosa* populations studied. NT = Niedere Tauern (Austria), FG = Făgăraș (Romania), VT = Vysoké Tatry (Slovakia), ZT = Západné Tatry (Slovakia), colored by ecotype (black = foothill ecotype, grey = alpine). We estimated the average values of temperature and solar irradiance (Photosynthetic Active Radiation [PAR]) over April, May and June that corresponds to the growth period of *A. arenosa*. The two climatic variables were obtained from the high-resolution climate database SolarGIS, version 1.9, operated by GeoModel Solar (Bratislava, Slovakia). Briefly, air temperature was derived from the Climate Forecast System Reanalysis and Global Forecast System databases (© National Centers for Environmental Prediction, USA) at 2 m height over the period 1990 to 2009 using a 1 km resolution and PAR from satellite and atmospheric data resolution over the period 2003 to 2013 using a 500 m resolution. B) Schematic representation of the four pairs of foothill and alpine ecotype and illustrative photos of typical foothill and alpine habitat.

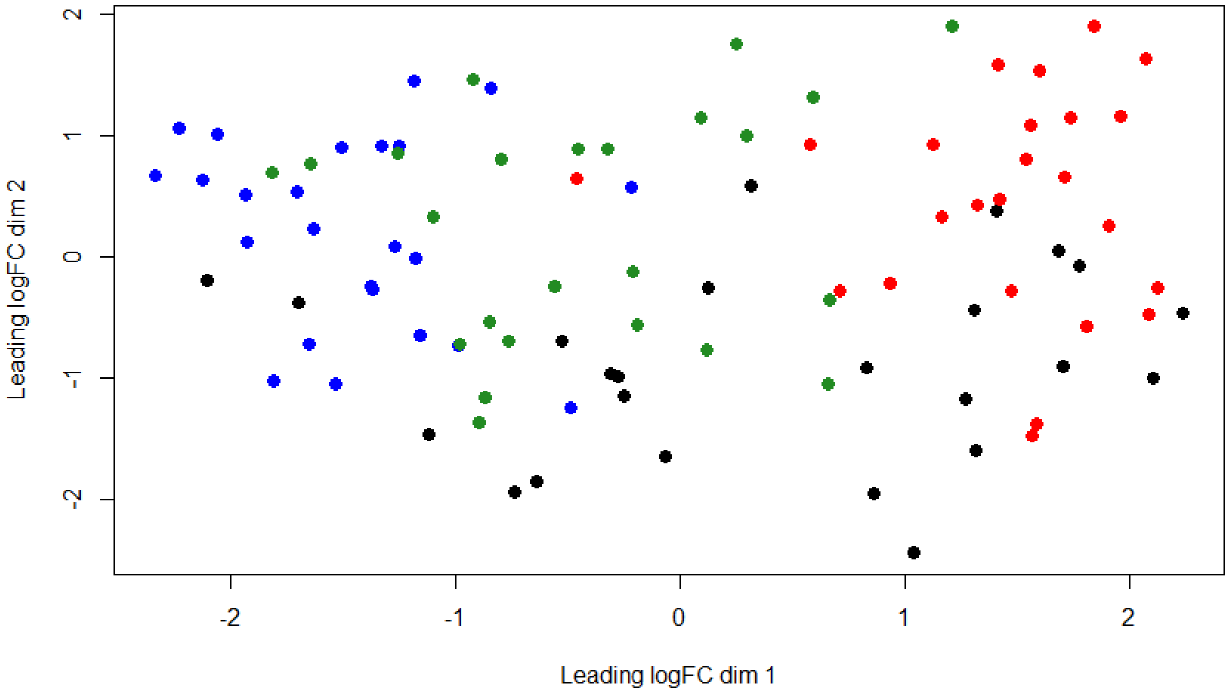


**Fig. S2.** Multidimensional scaling (MDS) plot showing expression differences between the 96 samples coloured by treatment: High T°C:High irradiance in red, High T°C:Low irradiance in black, Low T°C:High irradiance in green and Low T°C:Low irradiance in blue.

**Table S8**. Number of parallel outlier SNPs (SNPs identified as differentiation outliers in both compared regions) identified in our set of parallel DEGs compared to the total number of parallel outlier SNPs within the same pair of regions. The last column shows p-value of the enrichment analysis using hypergeometric test. NT = Niedere Tauern (Austria), FG = Făgăraș (Romania), VT = Vysoké Tatry (Slovakia), ZT = Západné Tatry (Slovakia).

| **Pairs of regions** | **Cis-regulatory elements** | | | **Coding elements** | | |
|---|---|---|---|---|---|---|
| | **N of parallel outlier SNPs identified in parallel DEGs** | **Total N of parallel outlier SNPs** | **Enrichment** | **N of parallel outlier SNPs identified in parallel DEGs** | **Total N of parallel outlier SNPs** | **Enrichment** |
| NT-FG | 5 | 519 | 0.390 | 2 | 366 | 0.859 |
| NT-VT | 7 | 992 | **0.041*** | 0 | 570 | 0.959 |
| NT-ZT | 11 | 968 | 0.289 | 4 | 572 | 0.869 |
| FG-VT | 43 | 3052 | 0.439 | 31 | 2348 | 0.652 |
| FG-ZT | 73 | 3062 | 1 | 88 | 2416 | **0.004**** |
| VT-ZT | 176 | 11294 | 0.208 | 99 | 7088 | 0.827 |
